# Segway 2.0: Gaussian mixture models and minibatch training

**DOI:** 10.1101/147470

**Authors:** Rachel C.W. Chan, Maxwell W. Libbrecht, Eric G. Roberts, William Stafford Noble, Michael M. Hoffman

## Abstract

**Summary:** Segway performs semi-automated genome annotation, discovering joint patterns across multiple genomic signal datasets. We discuss a major new version of Segway and highlight its ability to model data with substantially greater accuracy. Major enhancements in Segway 2.0 include the ability to model data with a mixture of Gaussians, enabling capture of arbitrarily complex signal distributions, and minibatch training, leading to better learned parameters.

**Availability and Implementation:** Segway and its source code are freely available for download at https://segway.hoffmanlab.org. We have made available scripts (https://doi.org/10.5281/zenodo.802940) and datasets (https://doi.org/10.5281/zenodo.802907) for this paper’s analysis.

## 1 Introduction

Segway identifies recurring combinatorial patterns in multiple genome-wide signal datasets such as ChIP-seq or DNase-seq data (Hoffman *et al*., 2012). Segway uses discovered patterns to assign a label to every position in the genome, resulting in a semi-automated genome annotation. It is commonly used to define chromatin state across the whole genome by resources such as the ENCODE Project (ENCODE Project Consortium, 2012) or the Ensembl Regulatory Build (Zerbino *et al*., 2015). Using chromatin data, the labels might represent genomic features such as “enhancer” or “facultative heterochromatin”.

Since its initial publication, we have made many changes to Segway. Release notes (https://bitbucket.org/hoffmanlab/segway/src/default/NEWS) contain a complete list. Of the new features, we expect the new standalone mode to interest the most users. This mode removes the requirement for a cluster system such as Sun Grid Engine, allowing one to run Segway easily on any Linux host. Also of interest are new features which improve Segway’s ability to learn more complex patterns with less configuration. Below we describe these features and demonstrate the improvement they provide.

## 2 Results

### Minibatch training

Segway uses the expectation-maximization (EM) algorithm to train its statistical model. Segway previously allowed for training only on a fixed region of the genome, such that each iteration of expectation-maximization (EM) training uses the same fixed region.

Using minibatch learning, each EM training iteration can now train on a different random region of the genome. For example, using a minibatch fraction of 1%, each training iteration will now use a different random 1% of the genome. This eliminates concerns of overfitting to a fixed region, but since there is no longer any guarantee of convergence, the final set of emission parameters is chosen by evaluating the likelihood on a held-out validation set. In general, minibatch allows one to sample the whole genome without having to use the whole genome as the training set, which would be much slower. Since the minibatch feature selects a different random region of the genome to train on per iteration, there is also a very high variation in likelihood progression between instances, though the overall likelihood progression is, however, generally more positive than that for a fixed region.

To demonstrate, we trained Segway on ENCODE (ENCODE Project Consortium, 2012) GRCh38/hg38 ChIP-seq datasets for H3K4me1 (ENCFF509XSM), H3K4me3 (ENCFF745GML), H3K27ac (ENCFF890NAY), H3K27me3(ENCFF592CSV) andCTCF(ENCFF884IIL) in the cell-line DOHH2 (Kluin-Nelemans *et al*., 1991). We did this using a single Gaussian model for a randomly selected fixed region of size 1%, and using minibatch training with batch size 1%. For each training round, we evaluated the posterior log likelihood of its learned parameters on 1.5% of the genome, which we held out from training in all cases. Minibatch resulted in a higher log likelihood convergence on the validation set both on average and in the final winning set of parameters (Figure 1). The fixed case also suffered from the validation set likelihood dropping from its initial peak due to overfitting on the training set (Figure 1).

**Fig. 1.**
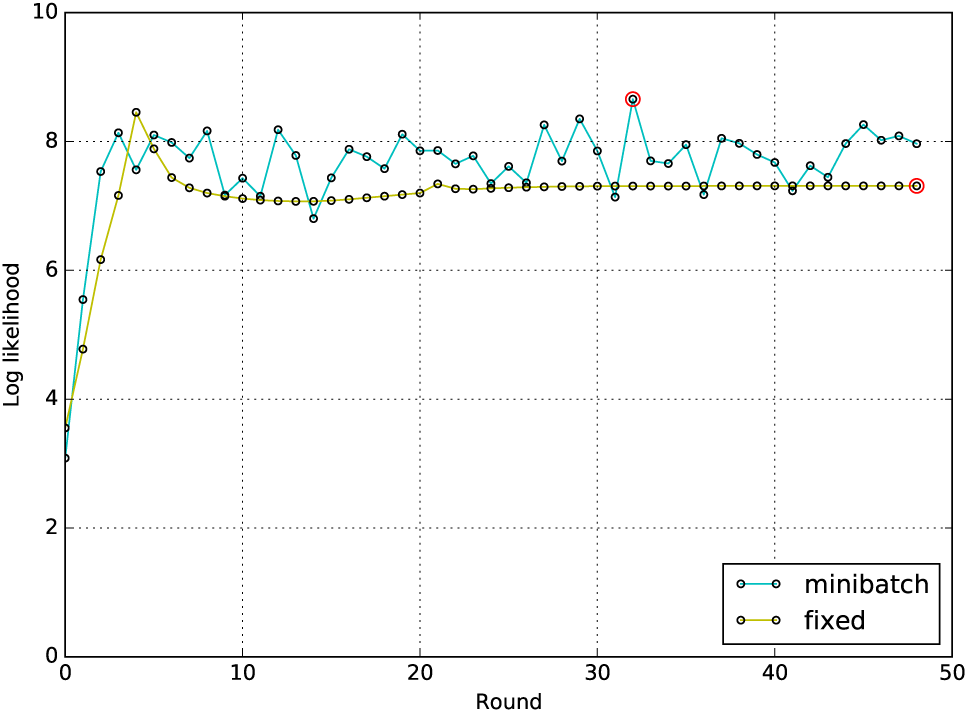
Log likelihood progression against round for a fixed 1% of the genome and a 1% chosen fraction for minibatch. Each series shared the same set of starting parameters and were evaluated against the same held-out validation set. Red circles: likelihood for the final chosen set of parameters in each series.

### Gaussian mixture models

Segway learns Gaussian distributions over signal values to represent different patterns. Previously, Segway used a single-component Gaussian to model the signal in each dataset given some label such that there is one learned mean parameter for each track-label pair, and one fixed variance for a given track. To enable more complex signal distributions, we extended Segway’s model to allow for a mixture model with k Gaussian components. Now, there are k mean parameters for each track-label pair, and k variances for each track. Using a mixture of Gaussians allows learning emission distributions that can more accurately fit data distributed non-normally.

To demonstrate, we trained Segway on signal data for the histone mark H3K27ac in the cell line DOHH2, using a 1-component Gaussian model and using a 3-component mixture of Gaussians. As previously described (Roberts *et al*., 2016), we trained using minibatch on 1% of the genome, for 10 labels and 100 EM training iterations. For each label learned, we extracted all datapoints corresponding to that label in the final annotation to generate an empirical distribution. Using the Kolmogorov-Smirnov test statistic, *D*, we picked the labels that best fit the empirical distribution in each case and compared them (Figure 2). The smaller the *D* statistic, the closer the fit between the theoretical and empirical distributions. Because Segway performs unsupervised learning, the sets of labels between each case do not correspond identically.

**Fig. 2.**
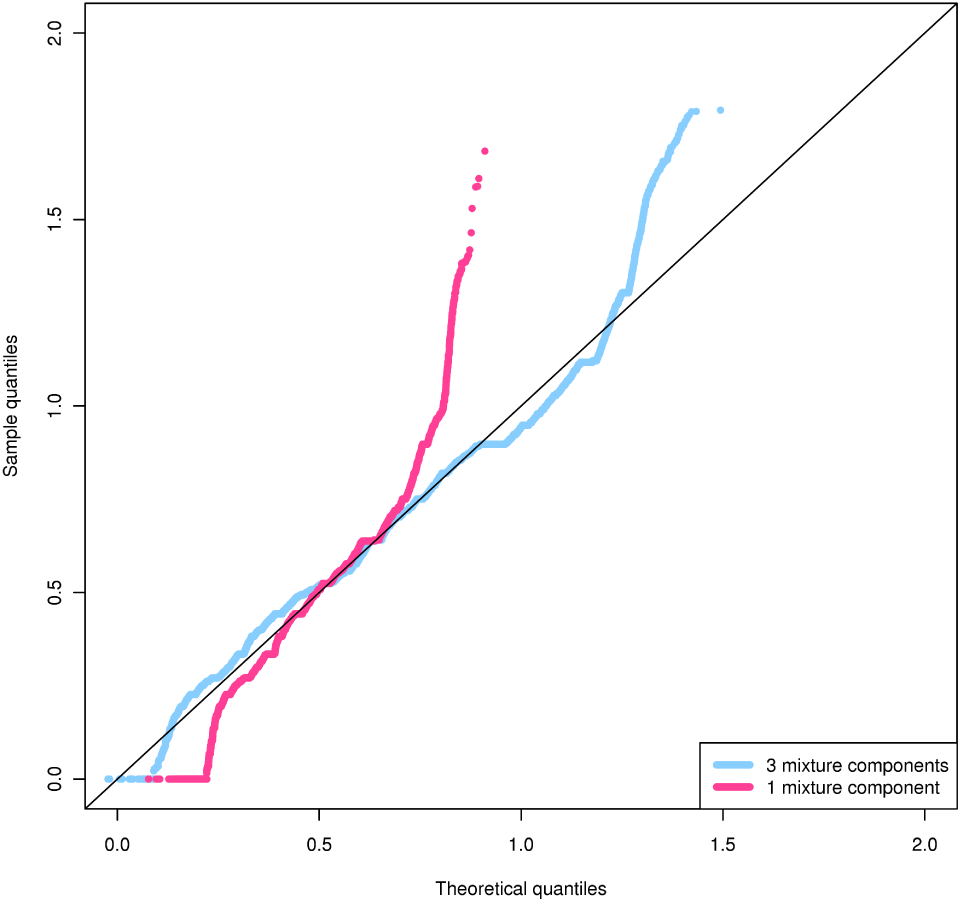
Combined quantile-quantile plot demonstrating ability of 1 or 3 Gaussian components to capture their empirical distributions.

In the single-component Gaussian case, the average *D* statistic across all labels was 0.28, with a median of 0.29, and a best *D* statistic of 0.078. In the 3-component mixture of Gaussians case, the average *D* statistic across all labels was 0.16, with a median of 0.10, and a best *D* statistic of 0.058.

The theoretical distribution for the 3-component mixture of Gaussians agreed with its multi-modal empirical distribution except for a slight right-skew in the data (Figures 2, S1). In comparison, the theoretical distribution for the single-component Gaussian model does not agree very well with its empirical distribution, with a strong skew in the tails of the distribution (Figure 2).

In conclusion, the mixture of Gaussians model better captures the empirical distribution than the single-component Gaussian model both on average and overall.

## Acknowledgments

This work was supported by the Natural Sciences and Engineering Research Council of Canada (RGPIN-2015-03948 to M.M.H.), the Canadian Institutes of Health Research (384410 to R.C.W.C.), and the National Institutes of Health (U41HG007000 to W.S.N.).

## Supplemental figures

**Fig. S1.**
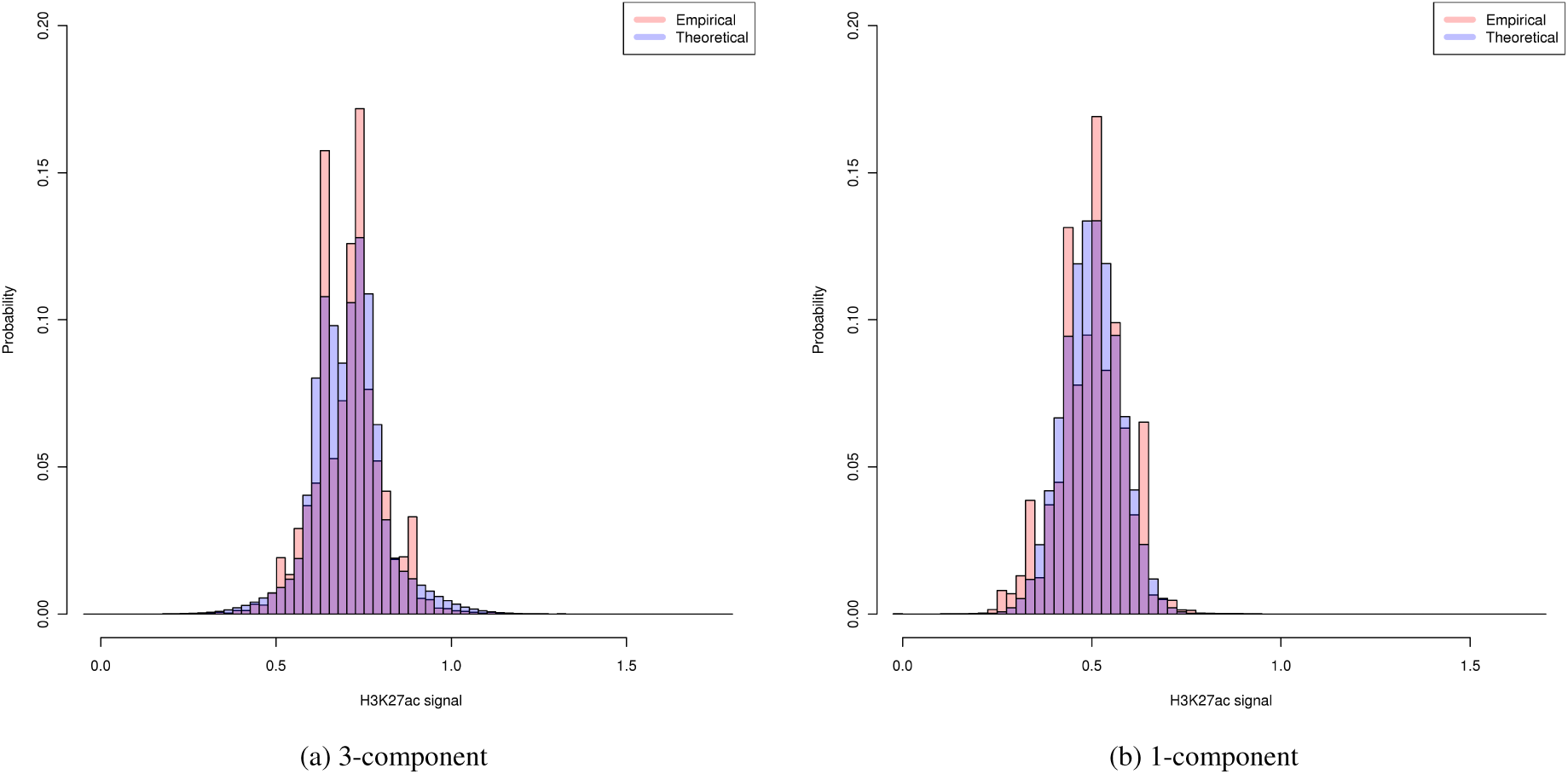
Histograms for the 3 (left) and 1 (right)-component mixtures of Gaussians showing the labels with the best *D* statistic in each case. The histograms are between the datapoints underneath that label (pink bins) against the same number of datapoints drawn randomly from the theoretical distribution of that label (blue bins).

